# Kinetics and topology of DNA associated with circulating extracellular vesicles released during exercise

**DOI:** 10.1101/2021.02.12.430930

**Authors:** Elmo W. I. Neuberger, Barlo Hillen, Katharina Mayr, Perikles Simon, Eva-Maria Krämer-Albers, Alexandra Brahmer

## Abstract

Although it is widely accepted that cancer derived extracellular vesicles (EVs) carry DNA cargo, the association of cell-free circulating DNA (cfDNA) and EVs in plasma of healthy humans remains elusive. Using a physiological exercise model, where EVs and cfDNA are synchronously released, we aimed to characterize the kinetics and localization of DNA associated with EVs. EVs were separated from human plasma using size exclusion chromatography or immuno-affinity capture for CD9^+^, CD63^+^, and CD81^+^ EVs. DNA was quantified with an ultra-sensitive qPCR assay targeting repetitive LINE elements, with or without DNase digestion. This model shows that a minute part of circulating cell-free DNA is associated with EVs. During rest and following exercise, only 0.12 % of the total cfDNA occurs in association with CD9^+^/CD63^+^/CD81^+^EVs. DNase digestion experiments indicate that the largest part of EV associated DNA is sensitive to DNase digestion and only ~20 % are protected within the lumen of the separated EVs. A single bout of running or cycling exercise increases the levels of EVs, cfDNA, and EV associated DNA. While EV surface DNA is increasing, DNAse-resistant DNA remains at resting levels, indicating that EVs released during exercise (ExerVs) do not contain DNA. Consequently, DNA is largely associated with the outer surface of circulating EVs. ExerVs recruit cfDNA to their corona, but do not carry DNA in their lumen.

## 1. Introduction

All cells of the human body are constantly and actively releasing a large amount of molecular material into the extracellular space. Directly or indirectly those extracellular molecules can interact in signaling pathways having crucial roles in the development of homeostasis and the coordination of physiological processes [1]. Acute physical exercise is a relevant stressor to disrupt homeostasis and trigger the release of a plethora of molecules into the circulation. A single bout of exercise affects thousands of molecules which orchestrate biological processes including energy metabolism, oxidative stress, inflammation, growth factor response, as well as other regulatory pathways [2]. Next to the release of proteins, collectively referred as secretome [3], nucleic acids including DNA [4–6], and RNA [7] are released under resting conditions and increase during exercise [2,8–10]. Circulating cell-free DNA (cfDNA) is mainly released from cells of the hematopoietic lineage, at rest [11,12], and during exercise [13]. cfDNA shows a typical fragmentation pattern, with a peak at ~166 bp representing a stretch of DNA wrapped around a nucleosome connected to histone 1, which protects DNA from nuclease degradation [14]. Additionally, it has been suggested that cfDNA is associated with EVs to be protected from degradation [1,15].

EVs are versatile mediators of cellular communication found in various body fluids, including blood [16], where their abundance also increases in response to acute physical exercise (termed ExerVs) [17–19]. In human blood a heterogeneous mixture of EVs is present [20,21]. They can be classified due to their cellular origin into microvesicles (MVs, 100-1000 nm in size), which shed from the outer plasma membrane, exosomes (30-100 nm) with endosomal origin, and apoptotic bodies, which are produced in the process of apoptosis and range between 1-5 μm in diameter, though smaller vesicles < 1000 nm are also released [16]. EVs compose of a phospholipid bilayer membrane that enables protected transport of bioactive cargo like proteins, lipids and nucleic acids between cells and tissues inside the vesicle. Additionally, macromolecules can be attached outside of the EV as part of the vesicle corona [22,23].

Although recent research indicates that nuclear DNA is a cargo of EVs in cancer, the association of DNA and EVs in healthy plasma is incompletely understood and controversially discussed (reviewed in [1,15,24]). A number of studies determined the association in cell culture and tumor models [25–34], as well as human plasma or serum of cancer patients [35–40], and healthy individuals [26,38,39,41–43]. In studies analyzing serum and plasma, an EV-associated part of the cell-free DNA of up to 90 % has been reported [36,43], whereas others find only a minor part associated to EVs [37,41], or no DNA in healthy individuals [39]. These differing observations are likely influenced by the heterogeneity and the different origin of EVs (cancer patient versus healthy person versus cell culture), as well as technical disparities of EV isolation and DNA quantification methods.

Separation of EVs from cell culture media or blood plasma is commonly performed using ultracentrifugation (UC) approaches, polymer-based precipitation, size exclusion chromatography (SEC), or immuno-precipitation for markers which are present on the vesicle membrane (reviewed in [44]). Isolation by differential UC (dUC), polymer-based precipitation as well as SEC lead to the co-isolation of high amounts of plasma proteins and lipoproteins [45]. EV preparations that include density gradient centrifugation using sucrose or iodixanol lead to high purity and further allows a discrimination of EV subpopulations, but are laborious and require high sample input [46]. A quick method to separate EVs from plasma with reduced contamination of plasma proteins and lipoproteins is immuno-affinity capture of EVs using magnetic beads coupled to antibodies for the tetraspanins CD9, CD63, and CD81 [47]. Still, currently no isolation method exists, that purifies only one EV subtype free of co-isolated non-EV material (protein aggregates, lipoproteins, etc.), which is especially important when studying EVs and their cargo and functions in body fluids [45,48].

Here, we elucidate the relationship between cfDNA and EVs under physiological conditions and following physical exercise. We separate EVs from plasma of exercising humans, implementing two different EV preparation methods including SEC, and immuno-affinity capture for CD9, CD63, and CD81. Via ultra-sensitive qPCR, directed against the repetitive LINE-1 element, DNA amounts in plasma, EV-depleted plasma, and isolated EVs are determined. Finally, by using DNase I and proteinase K digestion we study the amount of DNA, which is associated with the EV surface, or enclosed into the vesicular lumen.

## 2. Materials and Methods

### 2.1. Ethics Approval

Healthy human subjects were recruited at the Department of Sports Medicine, Johannes Gutenberg-University Mainz, Germany. The experimental procedures were approved by the Human Ethics Committee Rhineland-Palatinate and adhere to the standards of the Declaration of Helsinki of the World Medical Association. All subjects were informed about the procedures and the aim of the study and gave written consent to participate.

### 2.2. Subjects and Exercise Testing

A total of 10 healthy subjects participated in the study. Exclusion criteria were any diseases or signs of infections, as well as the use of prescribed medications, including anticoagulant treatment. Exercise testing was performed in the morning after a minimum of 8 h over-night fasting. Five of the subjects conducted an all-out incremental exercise test on a treadmill. One week later the same subjects conducted an all-out incremental exercise test on a bicycle ergometer. A second cohort of five subjects did an all-out incremental exercise test on a treadmill only. For the treadmill tests the participants started at a speed between 4-6 km/h, according to their expected fitness level. Every three minutes the speed was increased for 2 km/h with 30-45 sec break between each increment. Cycling tests started at 40 Watt. Every three minutes the load was increased for 40 Watt. The participants stopped the tests volitionally after exhaustion. After each step the subjects were asked for their rating of perceived exhaustion (RPE), values between 6-20 were possible [49]. Heart rate (HR) was measured continuously.

### 2.3. Blood Sample Collection and Plasma Preparation

At rest (Pre), immediately after (Post), and 30’ after the exercise bout venous blood was collected from the median cubital vein with a Safety-Multifly needle (0.8 × 19 mm) (Sarstedt) and collected in tripotassium-EDTA covered 7.5 ml Monovettes (Sarstedt). Platelet-free plasma was prepared within 5 min after blood drawing by two rounds of centrifugation for 15 min at 2,500 x g at room temperature [50]. Plasma was aliquoted and kept on ice or +4°C until EV isolation, to avoid any freezing of the samples. Samples that were used for estimation of total cfDNA in plasma were stored at −80°C until measurement.

### 2.4. Extracellular Vesicle Isolation

EVs were purified from plasma either by SEC or immuno-affinity capture followed by magnetic separation. SEC was performed as described in Brahmer et al., 2019 [17]. Briefly, 2 ml of plasma were layered on a self-made SEC-column (10 ml column volume, Sepharose CL-2B, Sigma-Aldrich) and a maximum of 24 one ml-fractions were collected by constantly adding PBS (Figure 1a).

**Figure 1.**
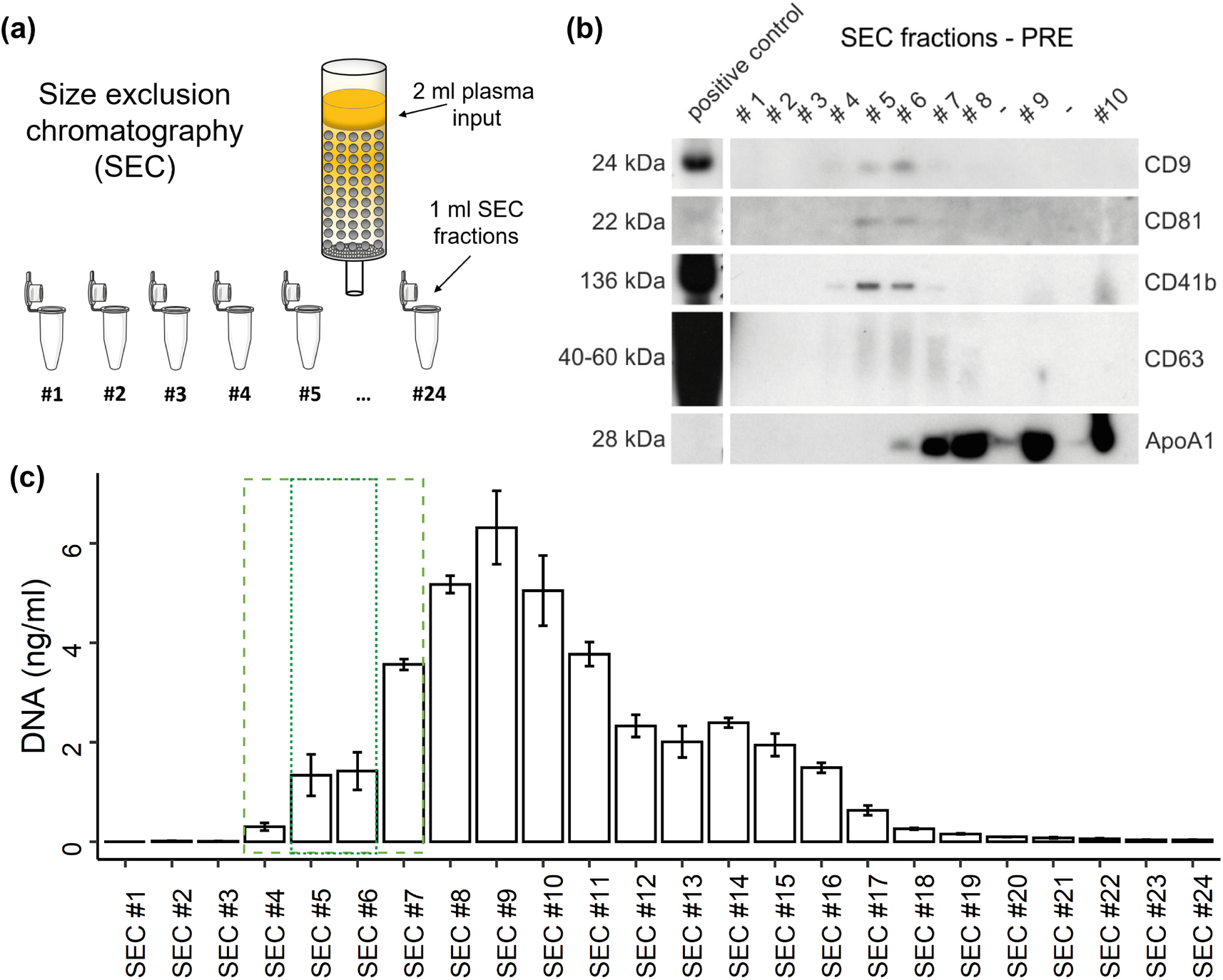
Distribution of EVs and DNA in SEC fractions. **(a)** Illustration of size exclusion chromatography (SEC) of plasma. **(b)** Western blot (WB) analysis of specific SEC fractions using genuine EV-markers including CD9, CD81, and CD63, as well as platelet specific marker CD41b, and lipoprotein marker ApoA1. **(c)** Concentration of DNA in SEC fractions measured by qPCR. The dashed box indicates SEC fractions containing EVs. The dotted box highlights most EV rich fractions.

**Figure 2.**
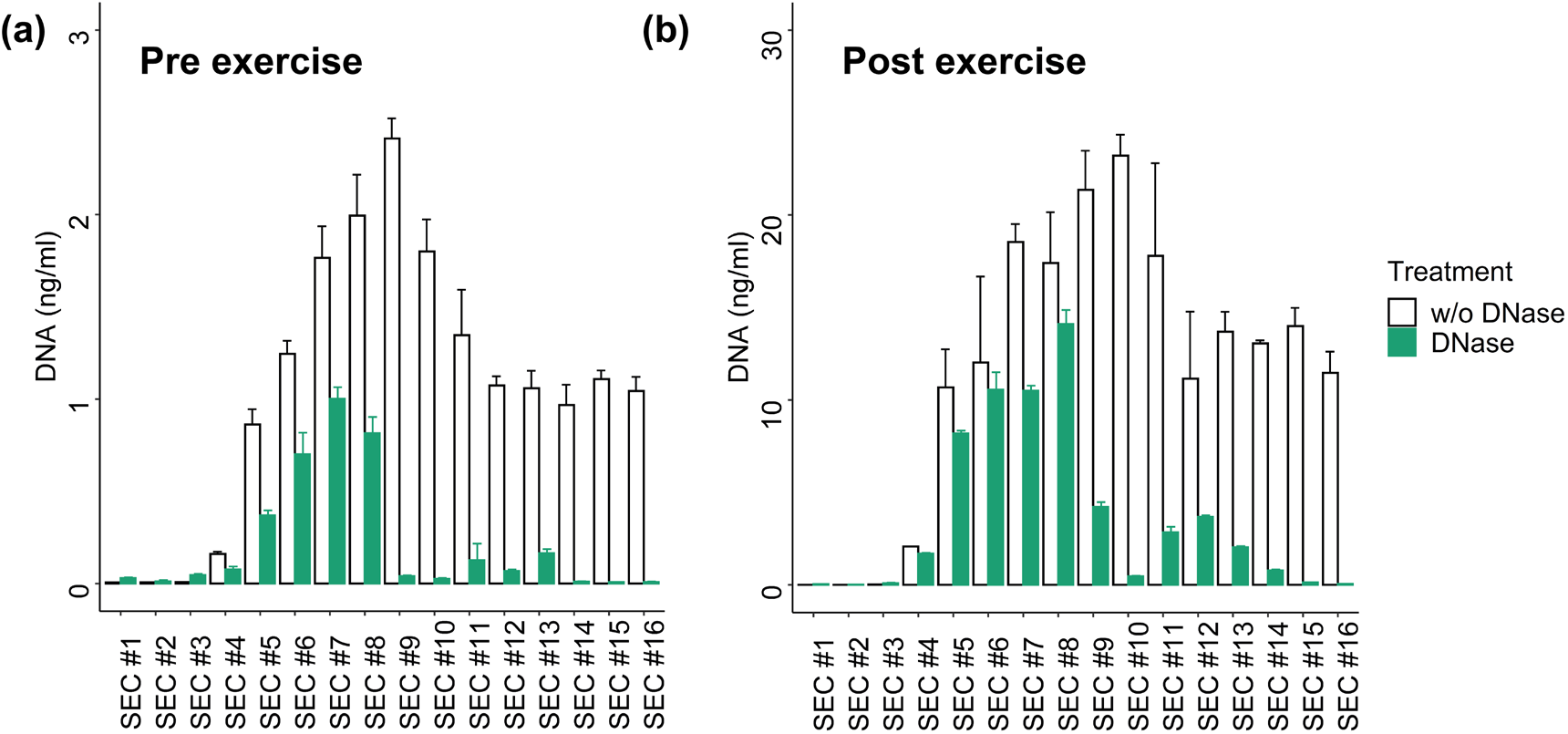
Distribution of DNA in SEC fractions with or without DNase I digestion of (a) Pre exercise plasma sample and (b) Post exercise sample. The DNA concentration was measured with qPCR without prior DNA isolation. Post exercise values show ~10 fold higher DNA concentrations.

For immuno-affinity capture, the Exosome Isolation Kit Pan, human (Miltenyi Biotec) was used according to the manufacturer’s instructions. Briefly, 50 μl of a mixture of anti-CD9, anti-CD63, and anti-CD81 magnetic beads were added to 2 ml of plasma and incubated for 1 h with constant shaking. Subsequently, EVs were magnetically captured, washed and eluted in a final volume of 100 μl. Immuno-affinity captured CD9^+^/CD63^+^/CD81^+^EVs as well as EV-rich and EV-poor SEC fractions were aliquoted and either prepared for western blotting or treated for cfDNA isolation and/or measurement as described below. One subject was excluded from the analysis since EV purification was markedly impaired: The magnetic bead isolation of the post sample repeatedly showed low bead recovery, indicated by the color of the eluate. Additionally, the SEC column clogged during separation.

We have submitted all relevant data of our experiments to the EV-TRACK knowledgebase (EV-TRACK ID: EV210058) [51].

### 2.5. Treatment of SEC- or CD9^+^/CD63^+^/CD81^+^EV Samples and DNA Isolation

To study the distribution of cfDNA in the SEC samples and to analyze the susceptibility to DNase I digestion, 20 μl of SEC samples were either pre-treated for 40 min at 37 °C with DNase I (Roche) at a concentration of 1 IU/μl, with 10 x Reaction Buffer, or mock treated with PBS. 2 μl of the samples were used for direct measurement of DNA without prior DNA isolation.

To study the association of DNA and CD9^+^/CD63^+^/CD81^+^EVs as well as the proportional part of DNA which is inside the EVs, the isolated EVs were aliquoted in 20 μl fractions and treated with or without TritonX100 (TX100; CarlRoth), proteinase K (CarlRoth), and DNase I (Roche) as follows: The samples were preincubated with TX100 (0.5 % final) and/or proteinase K (50 μg/ml final) with CaCl2 (CarlRoth) at a final concentration of 5 mM in PBS (Sigma). All sample were incubated for 30 min at 37°C, shaking at 250 rpm. The proteinase K activity was subsequently inhibited by adding 5 mM phenylmethylsulfonyl fluoride (PMSF) at room temperature. DNaseI at a final concentration of 1 mg/ml and 10 x DNase reaction buffer were added and the samples were incubated for 40 min at 37 °C with shaking, before DNA isolation using the QIAamp DNA Micro Kit (Qiagen), according to the manufacture’s recommendations. Briefly, the samples were filled to 100 μl with buffer ATL, before adding 10 μl proteinase K, and 100 μl buffer AL. All samples were heat incubated at 56 °C for 10 min. 50 μl EtOH (CarlRoth) were added before the isolation of DNA using the silica columns. The samples were eluted in a final volume of 20 μl of H2O.

### 2.6. Western Blotting (WB)

CD9^+^/CD63^+^/CD81^+^EVs or SEC fractions were mixed with WB sample buffer (200 mM Tris-HCl (pH 6.8); 10 % SDS; 0.4 % bromophenol blue; 40 % glycerol; 400 mM DTT; non-reducing conditions for CD9 and CD63 antibodies) and heated for 10 min at 70 °C. Volume-normalized samples were subjected to SDS-PAGE (12 % gels) and WB using PVDF-membranes. Membranes were blocked (4 % milk powder and 0.1 % Tween in PBS) and incubated with primary and HRP-coupled secondary antibodies followed by chemiluminescent detection.

The following antibodies and dilutions were used: CD9 (1:2000 dilution, clone #MM2/57, Merck Millipore), CD63 (1:500 dilution, #CBL553, Merck Millipore), CD81 (1:1000 dilution, #B-11, Santa Cruz), CD41 (1:1000 dilution, #SZ.22, Santa Cruz), ApoA1 (1:200 dilution, #12C8, Santa Cruz), goat-anti-mouse-HRP (1:10,000 dilution, polyclonal, 115–035-166, Dianova).

### 2.7. Cell-free DNA Measurement

The cfDNA concentration was determined with a quantitative real-time PCR (qPCR) assay, targeting a hominoid specific 90 bp repetitive DNA element [52]. The repetitive element occurs 3416 times in the genome enabling very sensitive and reliable quantification. The primer sequences are 5’- TGCCGCAATAAACATACGTG-3’ and 5’- GACCCAGCCATCCCATTAC −3’ for the forward and reverse primer, respectively. The following cycling conditions were used with a CFX384 BioRad cycler: 2 min 98 °C heat activation, followed by 10 sec 95 °C and 10 sec 64 °C for 35 cycles and subsequent melting curve from 70 – 95 °C with 0.5 °C increments for 10 sec. Each sample was measured in triplicate of 5 μl with the following final concentrations: Velocity Polymerase 0.6 U (Bioline), 1.2 x Hifi Buffer (Bioline), 0.1 x SYBR Green (Sigma), 0.3 mM dNTPs (Bioline), 140 nM of each primer. 2 μl of sample were mixed with 13 μl of mastermix. The amount of DNA was calculated as described in Neuberger et al. [52], and is briefly described in Appendix A. Plasma as well as the flow through of the immuno-affinity isolations were diluted 1:10 in ultra-pure H2O (Invitrogen) or 1 x PBS (Sigma). To investigate if immuno-beads in the CD9^+^/CD63^+^/CD81^+^EV samples inhibit qPCR results, spike-in experiments were performed. Undiluted or diluted mock immuno-affinity isolates (1:1, 1:5, 1:10) and 1 x PBS were spiked with 200 or 20,000 copies of the L1PA2 target sequence. A dilution of 1:1 did not show any inhibitory effects and was used as the dilution of CD9+/CD63+/CD81+EV samples.

### 2.8. Data Analysis

The qPCR data was captured with the CFX Manager Software, Version 3.0 (Bio-Rad) and Microsoft^®^ Excel, 2016. Statistical analysis was conducted with R version 4.0.3, using tidyverse version 1.3.0, and rstatix version 0.6.0 packages. The ggplot2 package version 3.2.2 was used for graphical illustrations. Continuous data was tested for normal distribution with Shapiro-Wilk test, after log normalization. On a global level repeated measures ANOVAs, including sphericity test, or the non-parametric Friedman test were performed. A significant global test was followed by post-hoc t tests, or Wilcoxon rank-sum tests, for normally or non-normally distributed data, repsectively. P < 0.05 was considered statistical significant (* = P < 0.05, ** = P < 0.01, *** = P < 0.001). Pearson correlation test was applied to study associations between normally distributed data. Otherwise the non-parametric Spearman correlation test was used.

## 3. Results

### 3.1. Participants characteristics and exercise performance

Ten participants, including 9 male and 1 female (age: 26.8 ± 4.49 y, height: 181.65 ± 6.8 cm, weight: 76.33 ± 8.03 kg, BMI: 23.09 ± 1.61), underwent all-out exercise tests. The five subjects who performed a repeated exercise test showed a mean of 5.42 min shorter time until exhaustion for the running exercise compared to cycling exercise (P = 0.028, 95 % CI = −9.89 - −0.94), with no significant differences between maximal heart rate (P = 0.23), and similar rating of perceived exertion (see Table 1).

**Table 1:** Participants characteristics and exercise performance. Values are given in mean (± SD). BMI = body mass index, RPE = rating of perceived exertion (values between 6 and 20 are possible).

| Subjects | Age (y) | BMI (kg/m<sup>2</sup>) | Exercise | Time until exhaustion (min) | Maximal Heart rate (1/min) | RPE |
| --- | --- | --- | --- | --- | --- | --- |
| n=5 | 23.8 (± 1.47) | 22.95 (± 1.57) | Running | 22.96 (± 2.56) | 193.4 (± 2.87) | 19.6 (± 0.49) |
| | | | Cycling | 28.32 (± 3.54) | 188.2 (± 8.30) | 19.8 (± 0.40) |
| n=5 | 29.6 (± 4.72) | 23.56 (± 1.06) | Running | 21.10 (± 1.56) | 189.8 (± 7.08) | 18.8 (± 0.40) |

### 3.2. Distribution of EVs and cfDNA in SEC fractions

To study the distribution of plasma cfDNA in platelet poor plasma after SEC, undiluted SEC samples were used for qPCR quantification without DNA isolation. WB analysis (Figure 1b) consistently shows the strongest signals for the genuine EV markers CD9, CD81, and CD63 in SEC fractions #5 and #6. Similarly, the platelet-derived EV marker CD41b becomes most detectable in those fractions. Overall, EV markers are detectable in SEC fractions #4-7, whereas in fraction #7 already a strong ApoA1 signal appears, indicating a relevant lipoprotein co-isolation. DNA starts to incline in SEC #4, showing the highest concentration in SEC #9 (Figure 1c). The DNA concentrations display a plateau from SEC #12 to #15, declining sharply after SEC #16. About 17.2 % of the total DNA (38.5 ng) occurs in the SEC fractions #4-7 while a part of 7.17 % occurs in the fractions 5 and 6, which show highest EV amounts. The major part of cfDNA is found in non-EV SEC fractions and only a minor part can possibly be associated with EVs.

### 3.3. DNA digestion in SEC fractions

To get an impression about susceptibility of the DNA contained in the distinct SEC fractions for DNase I digestion, a pre-exercise (Pre) and a post-exercise (Post) plasma sample were subjected to SEC and treated with DNase I. The total plasma cfDNA concentration increased ~9-fold from 16.65 ng/ml (Pre) to 173.40 ng/ml (Post) during exercise. The amount of DNA recovered in SEC samples (sum of SEC #1-16) was 16.85 ng in Pre and 173.4 ng in Post. The distributions of the cfDNA concentrations over the different SEC samples were highly similar between the Pre and Post samples (r = 0.977; P < 0.001). In the vesicular fractions (SEC #4-7) 21.61 % (Pre) and 24.98 % (Post) of the total DNA were detected. In the EV-rich fractions (SEC #5 and #6) are 11.49 % (Pre) and 13.08 % (Post) of the total DNA. DNAse I treatment slightly decreased the amount of DNA in fractions #4-7 from 4.03 ng to 2.14 ng in the Pre samples, and from 43.32 to 30.91 ng in the Post samples. In the remaining SEC fractions, more than 80 % of the DNA was digested. These results indicate that ~70-80 % of the total plasma cfDNA in Pre and Post conditions is not protected from DNase digestion, while the remaining part might be protected either by EVs or co-isolated lipoproteins or plasma proteins.

### 3.4. DNA associated with CD9^+^/CD63^+^/CD81^+^EVs

Since EV preparation by SEC is highly susceptible to co-isolate large lipoproteins, which are associated with extracellular plasma DNA [53], we decided to further study the association of DNA with EVs prepared by immuno-affinity capture, which shows higher purity. Figure 3a illustrates capture and magnetic separation of CD9^+^/CD63^+^/CD81^+^EVs from plasma. WB analysis confirmed the abundance of EV markers (a representative WB is presented in Figure 3b). Figure 3c displays the amount of DNA in 1 ml of plasma, 1 ml of flow through (FT), or CD9^+^/CD63^+^/CD81^+^EVs corresponding to 1 ml of plasma. The samples were taken Pre, Post, and 30′ after exercise, from n = 5 subjects who performed two repeated exercise tests. Multiple comparison t tests did not show significant differences between plasma cfDNA and FT cfDNA in any of the time points (P values without adjustment for multiple comparisons are presented, see Figure 3c). Figure 3e shows the percentage of EV-associated DNA with the total amount of plasma DNA. Overall, 0.12 ± 0.05 % (median ± SEM) of the total plasma DNA is associated with CD9^+^/CD63^+^/CD81^+^EVs. Specifically, 0.12 ± 0.08 % in the Pre samples, 0.15 ± 0.14 % in the Post, and 0.07 ± 0.02 % in the +30’ samples. The significant and strong correlation between plasma and CD9^+^/CD63^+^/CD81^+^EV cfDNA values (Figure 3d) indicates that a higher amount of DNA occurs in association with EVs if plasma cfDNA levels are higher.

**Figure 3.**
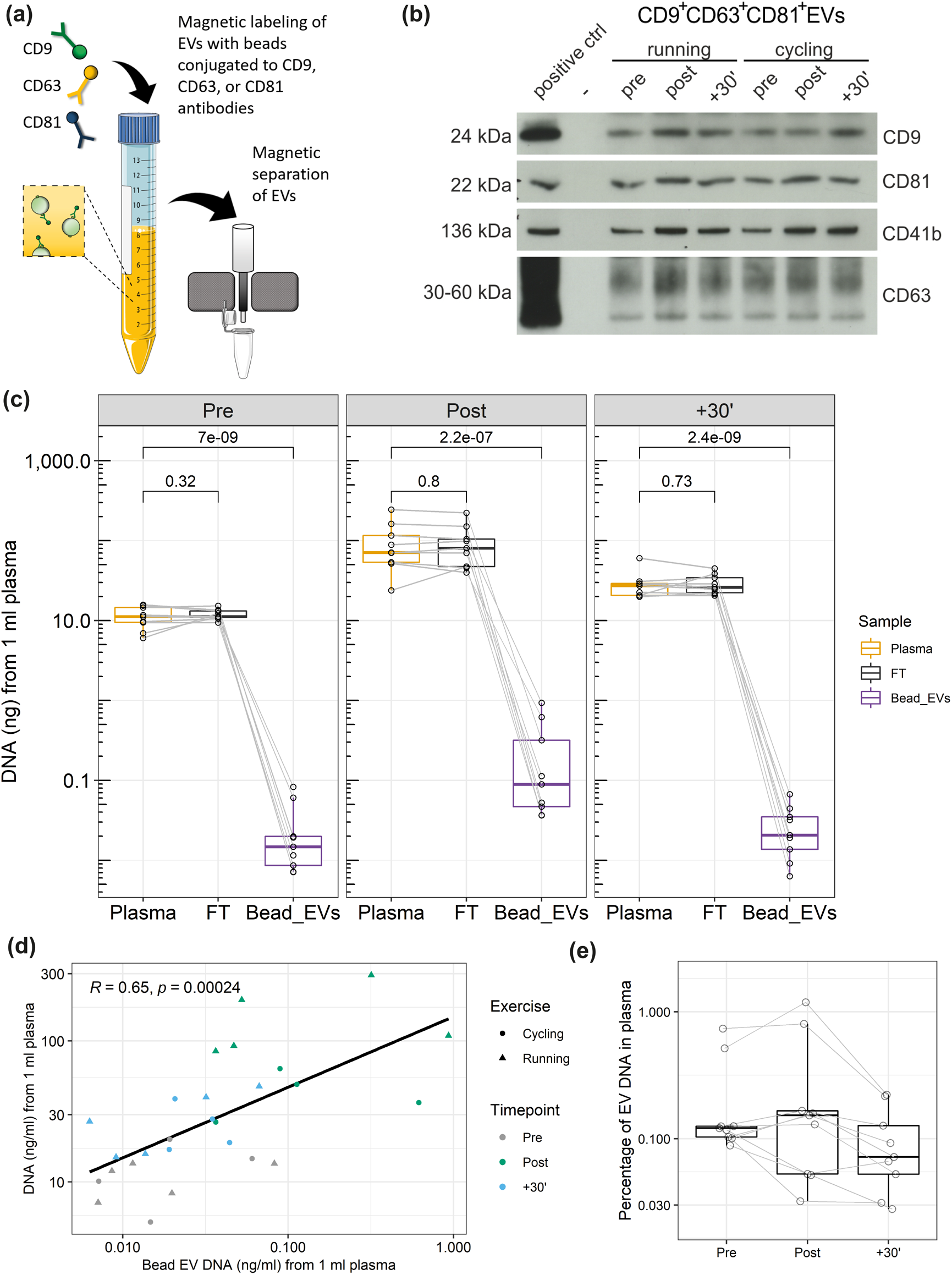
DNA associated to CD9^+^/CD63^+^/CD81^+^EVs. **(a)** Illustration of EV immunobead isolation (anti-CD9/CD63/CD81). **(b)** Western blot detection of EV proteins CD9, CD81, CD63, as well as platelet specific marker CD41b. **(c)** Amount of DNA in plasma, flow through, and CD9^+^/CD63^+^/CD81^+^EVs. The values represent the amount of DNA from 1 ml of plasma. **(d)** Correlation between DNA amount in plasma and DNA amount of CD9^+^/CD63^+^/CD81^+^EVs. **(d)** Percentage of CD9^+^/CD63^+^/CD81^+^EV DNA in relation to the total DNA amount in plasma.

Former studies indicated that different exercise modalities lead to different cfDNA increases. All-out treadmill running increased cfDNA levels ~10 fold [41,54], whereas all-out cycling increased plasma cfDNA only ~5 fold [17,55]. To evaluate the influence of the exercise setting on EV kinetics, five subjects performed each of the exercise tests. Figure 4 displays the fold changes of cfDNA in plasma and CD9^+^/CD63^+^/CD81^+^EV associated DNA, as well as WB results for CD9 and CD41b. Two-way repeated measures ANOVA indicated that plasma cfDNA levels differ significantly between time points and between exercise modalities running and cycling (F(2,8) = 14.55, P = 0.002). After cycling the cfDNA increased 4.06 fold. Running led to a 12.38 fold increase from Pre to Post (P = 0.013). The CD9^+^/CD63^+^/CD81^+^EV associated DNA increased significantly over time (F(2,6) = 28.67, P < 0.001), showing no differences between running and cycling (F(1,3) = 2.12, P = 0.242). Hence, in both cases the EV associated DNA increased similarly after running (fold change = 4.64) and cycling (fold change = 4.25). Likewise, for WB analysis of EV markers (CD9, CD41b, CD81) no significant difference was found between the exercise modalities, whereas the markers showed significant changes over time (CD9: F(2,8) = 13.04, P = 0.003; CD41b: F(2,8) = 15.38, P = 0.002; CD81: F(2,8) = 1.58, P = 0.021). The results confirm an increase of cfDNA and EVs after exercise. Notably, cfDNA concentration increases stronger after running compared to cycling, whereas no influence of the exercise setting was detectable on EV release and EV associated DNA, confirming independent release mechanisms [41].

**Figure 4.**
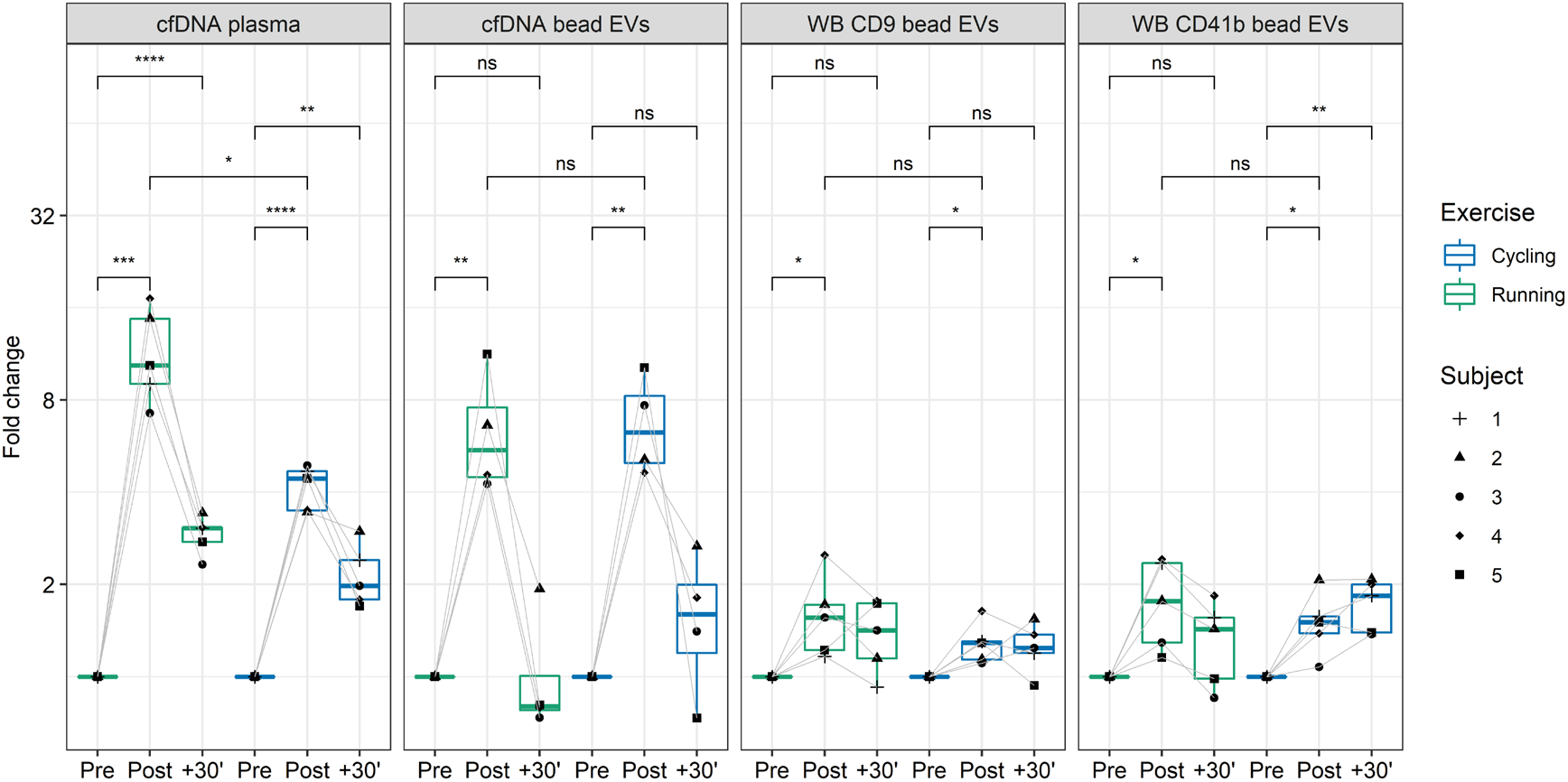
cfDNA and CD9^+^/CD63^+^/CD81^+^EVs kinetics in running versus cycling. The fold-changes of plasma cfDNA and CD9^+^/CD63^+^/CD81^+^EVs DNA, detected with qPCR, and fold changes of CD9^+^/CD63^+^/CD81^+^EVs, represented by CD9 and CD41b western blot analysis, in running and cycling exercise are illustrated. The labels represent selected comparisons between time points and exercise modalities, reflecting unadjusted p-values (* = P < 0.05, ** = P < 0.01, *** = P < 0.001).

### 3.5. DNA cargo of EVs

To study the proportion of DNA which is associated with the EV surface or protected within the lumen of EVs, CD9^+^/CD63^+^/CD81^+^EVs were treated as illustrated in Figure 5a, with or without DNase, and, in combination with protease or TX100. Subsequently, the DNA was isolated using a purification kit. DNase treatment should reduce the major part of DNA on the surface of EVs, however, a more efficient degradation could be reached in the presence of proteases that hydrolyse DNA-bound proteins [56]. A first comparison of DNase treated or untreated Pre and Post samples (Figure 5b), shows that the total amount of DNA increases significantly from Pre 9.51 (± 2.69) ng/ml to Post 77.37 (± 37.41) ng/ml (mean ± SEM). Notably, the DNase treated Pre and Post values do not differ significantly, showing similar DNA concentrations with Pre 3.61 (± 4.28) and Post 2.26 (± 1.50) ng/ml. This indicates that the cfDNA, which increased in response to exercise, is associated with the surface of EVs, susceptible for DNase digestion. As shown in Figure 5c and 5d, the DNase treatment reduced the amount of DNA in the Pre exercise samples from 9.51 (± 5.39) ng/ml (100 %) to 3.60 (± 2.14) ng/ml (43.88 %). DNase and protease reduced the concentration to 1.50 (± 0.39) ng/ml (20.11 %). TX100 and DNase treatment enabled the digestion of almost all DNA, reducing the value to 0.23 (± 0.02) ng/ml. The results indicates that ~20 % of the EV associated DNA are within the lumen of the EVs. The treatment with TX100 and protease without DNase showed similar results, compared to the untreated condition (7.85 ± 2.29 ng/ml, equaling 84.5 %). In the Post samples (Figure 5e and 5f) the DNase digest showed that almost all DNA is sensitive to DNase digestion, reducing the DNA concentration down to 4.47 % of the total EV associated DNA. Notably, in the Post samples the treatment condition protease with DNase showed slightly higher values than DNase alone (4.97 ± 1.12 and 2.26 ± 0.75 ng/ml, respectively). Additionally, in the TX100 and DNase treated samples remaining DNA was detected (3.27 ± 1.91 ng/ml), which might indicate that the DNase treatment did not work efficiently in the Post samples, which have about 8-fold higher DNA values compared to the Pre samples.

**Figure 5.**
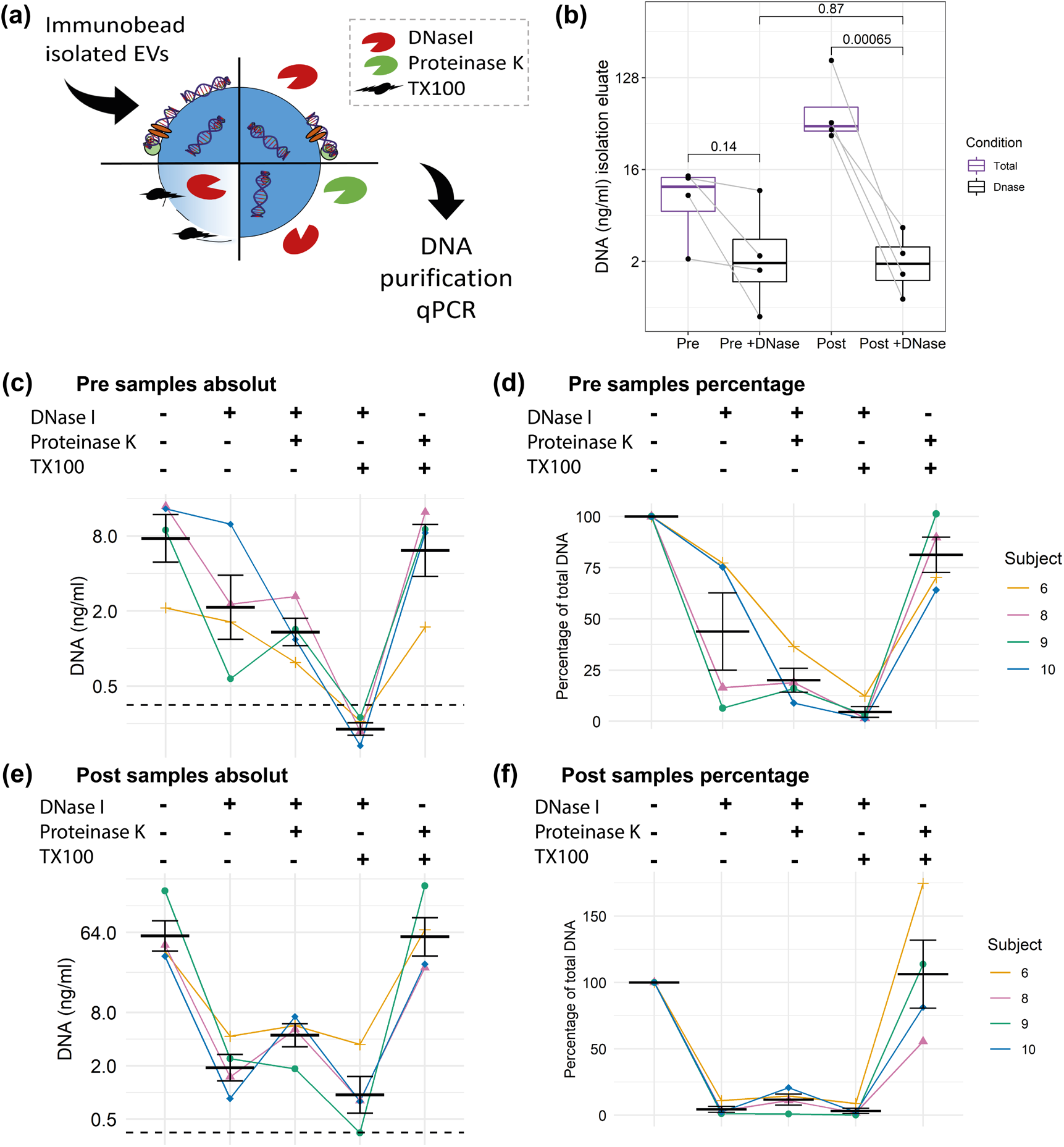
DNA cargo of CD9^+^/CD63^+^/CD81^+^EVs. (a) Illustration of different digestion strategies. (b) Comparison of Pre and Post samples with or without DNase I digestion. Unadjusted t test results are displayed. Absolut and relative concentration of the Pre samples (c,d) and Post samples (e,f). Error bars indicate mean ± SEM. The dashed black line indicates the limit of detection (LOD) of the qPCR.

## 4. Discussion

In spite of the great interest for cfDNA [57] and EVs [58] in the field of liquid biopsy, the association of the two entities has not been studied extensively [1,15,58]. Especially the association of DNA and EVs in the plasma of healthy human individuals remained elusive. Here we made use of acute exercise as a physiological model to study the concomitant release and relationship between EVs and cfDNA in the circulation. The results demonstrate that only a minute part of cfDNA is associated with EVs in human plasma. EV-associated DNA is largely associated with the surface as part of the corona and only found in trace amounts in the EV-lumen. Using a highly sensitive qPCR, we found a small amount of DNA in the lumen of immuno–affinity captured CD9^+^/CD63^+^/CD81^+^EVs, which was protected from DNase digestion. CfDNA and EVs both increase during exercise, whereas the cfDNA increase appears to occur completely independent of EVs and DNA is attached to the EV surface. Since the DNase-resistant fraction of EV-associated is constant from pre to post exercise, we conclude that the ExerV subpopulations do not contain luminal DNA.

We found that under physiological conditions only ~0.12 % of the total plasma DNA is associated with CD9^+^/CD63^+^/CD81^+^EVs (Figure 3e). Of this part ~20 % are protected within the lumen of the EVs indicated by DNAse digestion experiments (Figure 5). These results are in line with several previous findings on the association of DNA to EVs. In 2015, we studied the amount of extracellular DNA in plasma and EV sub-populations isolated by dUC [41], where most of the plasma DNA was detected in the supernatants after centrifugation. In 10.000x g pellets about 4.67 % of the DNA occurred, representing the DNA in larger MVs and apoptotic bodies. In the UC small EV pellet 3.94 % of the DNA occurred. An additional DNase digestion of the pellet led to further reduction to 0.94 %, which indicated that ~23 % of the EV associated DNA was protected from degradation. Similarly, Lázaro-Ibáñez et al., found DNA associated with low density and high density small EV populations prepared by density gradient centrifugation, which was mostly associated with the vesicle surface [37]. Only 20.9 % or 3 % of the DNA of TF-1 or HMC-1 cell EVs were protected from DNase digestion. Also, Fischer et al. found about one quarter of the DNA within the small EVs released from human mesenchymal stromal cells [28]. These results collectively indicate that only a minor part of cell-free DNA is associated to EVs and an even smaller part is actually transported encapsulated within the EVs.

A high amount of cfDNA cargo might be a characteristic of cancer cell derived EVs. A number of studies found DNA in cancer-derived EVs using different separation technologies including UC [36], dUC [25,40], density gradient centrifugation [37,39], flow field fractioning [32], as well as nano-flow cytometry [30]. Now it is widely accepted that DNA is a constituent of EVs in cancer patients [1], whereas DNA is more abundant in large EVs compared to small EVs [39]. More recent sub-characterization of the cancer small EVs into exosomal/non-exosomal [59], high density/low density populations [37], or distinct nanoparticles [33] / exomers [32], further challenge a clear association of DNA in small EVs. Zhang et al. found DNA associated with small EVs and distinct nanoparticles [33], whereas Jeppesen et al. did not find DNA in small EVs after DNase digestion [26]. Notably, cancer cells have a disrupted homeostasis and a direct comparison between so-called oncosomes and EVs released from healthy cells is hampered [39,40]. Our findings on a very small amount of DNA associated to EVs in healthy individuals are in line with the hypothesis that oncosomes are highly dissimilar to EVs in a physiological state.

Next to the cellular origin (cancer vs. healthy) the separation of EVs, as well as quantification of DNA will affect study outcomes. Contrasting our results, Fernando et al. described that up to 90 % of the plasma DNA is associated with EVs [43]. These differing results could be due to the choice of chemical precipitation as EV purification method. Precipitation based EV separation methods co-isolate a relevant amount of plasma macrocomplexes including lipoproteins [60,61]. The latter are likewise described to be carriers of extracellular DNA. About 12 % of the extracellular plasma DNA were shown to be associated to circulating lipoprotein complexes [53]. Hence, the quantified DNA amount in EV precipitates may overestimate the actual amount of EV-associated DNA. Notably, in our experiments using SEC for EV preparation not only the fractions showing EV markers (#4-#7) are protected from DNase digestion, but also other fractions are less sensitive to degradation (e.g. #8/#9, Figure 2). This might be a result of DNA association with other structures including lipoproteins or plasma proteins [62], highlighting the influence of the method of EV separation from plasma on experimental outcome.

Also, the method of cfDNA analysis is of high importance to study the DNA cargo of EVs. In a well-designed study, Vagner et al. analyzed the amount of DNA in dUC separated plasma EVs of 40 cancer patients and a subset of 6 healthy controls. The researchers detected 6-7 fold more DNA in large EVs, compared to small EVs of cancer patients. However, no DNA was detected in EVs separated from 1 ml of plasma from healthy subjects [39]. Notably, the High Sensitivity (HS) dsDNA Qubit Assay kit was used for DNA detection. During the handling process the samples are diluted at least 10 times in Qubit^®^ working solution [63]. The lower detection range of the kit is 1 or at least 0.5 ng/ml, whereas the values refer to the concentration of the diluted sample. Therefore, the sensitivity of the measurement device might not be sufficient, to detect the low concentrations of DNA associated with EVs. In contrast, our DNA quantification relies on a qPCR which amplifies repetitive DNA elements showing a very high sensitivity [52], which enables to measure the low DNA concentrations present in EVs.

Despite a similar cellular origin of EVs and cfDNA in blood following exercise suggests a joint release mechanism, we found further evidence for their independent release. Early sex-mismatch transplantation models showed that the major part of cfDNA is released from cells of the hematopoietic lineage [12]. In a similar study design we showed that hematopoietic cells are the major source of cfDNA during exercise [13]. More recently, the analysis of the methylation profile of cfDNA shows that about 32 % of cfDNA is derived from granulocytes, 30 % from erythrocyte progenitors, 23 % from monocytes and lymphocytes, 9 % from endothelial cells, and only 6 % from other cells including neurons and hepatocytes [11], while a sub-characterization for exercise released cfDNA has not been conducted yet. We and others found that next to muscle cells, platelets, endothelial cells, and leukocytes, significantly contribute to the pool of EVs released into plasma following physical exercise (reviewed in [48,64–66]). Like for cfDNA, the underlying signaling mechanisms are not fully understood, but an association with shear stress and the activation of coagulative processes are discussed [48]. Since cfDNA and EVs seem to be released by similar cell types during physical exercise, a joint or related release mechanism is possible, but remains speculative.

Intriguingly, our results indicate that cfDNA and EVs are not released as a single entity during exercise but likely associate with each other after their release. This assumption is based on two findings. First, as expected, the total cfDNA levels increased with a higher fold change after running exercise compared to cycling exercise (Figure 4). In contrast, the increase in the amount of DNA associated with CD9^+^/CD63^+^/CD81^+^EVs was similar (Figure 4). If cfDNA release would occur together with EVs, similar fold changes would have been expected. Secondly, after DNase digestion, Pre and Post exercise EV samples (Figure 5b) include a similar amount of DNAse-resistant DNA, although the EV-levels increase through the release of ExerVs. Indicating, that ExerVs do not contain DNA. Interestingly, the amount of surface bound DNA increased significantly during exercise, suggesting that cfDNA after its release is recruited to the EV surface and attached to the corona. As reviewed by Buzás et al., the EV corona can be physiologically relevant [23].

Overall, our study describes that only a minor part of cfDNA is associated with EVs in healthy humans. Our findings contrast with a high DNA association with oncosomes indicating distinct release mechanisms for the EV subtypes of different origin. It is conceivable that the minor and stable fraction of DNA-containing EVs in human plasma observed in the exercise paradigm reflect the steady state population of apoptotic bodies in the circulation, expected to contain fragments of DNA and known to remain constant within the timeframe of acute exercise. Furthermore, ExerVs appear to be free of luminal DNA. However, a single bout of all-out exercise significantly increases the amount of DNA bound to the surface of EVs. Surface DNA may well be of physiological significance for the adaptational processes induced by regular physical exercise, requiring further investigations. It will be interesting to examine the specific characteristics of surface-associated and luminal DNA of plasma EVs. Since we amplified a repetitive element which is distributed throughout the human genome, further research should emphasize the sequencing of the DNA. Despite only a minor part of DNA is EV associated, studying intraluminal and surface-bound DNA could reveal valuable information for the field of liquid biopsy linking EV DNA with physiological properties such as inflammation.

## Author Contributions

EMKA, PS, AB, EN conceived the original idea. BH, EMKA, PS, EN, AB conceived and planned the experiments. EN, AB, BH, KM performed experiments. EN and AB wrote the manuscript in consultation with EMKA, PS, BH and KM.

## Appendix A

To calculate the amount of DNA we utilized a validated qPCR assay [52]. Ahead of the measurements, the linearity, limit of quantification, limit of detection of the assay were established with three independent standard curves. In addition, a set of two reference samples were validated and are included in each run. The reference samples allow a calibration to account for inter-plate differences. To calculate the number of molecules in a 5μl qPCR the following formula is applied: 10^(cq – intercept)/slope^. The intercept and slope values are derived from the validated standard curves. A devision by 5 results in the number of molecules per μl. To calculate the number of genome equivalents per ml (GE/ml) the number of molecules are multiplied with the dilution factor of the sample and divided by 3416 (the number of hits of the amplified target in the human genome). The resulting genome equivalents are multiplied with 3.23 pg, the weight of a haploid genome. The full formula is as follows:

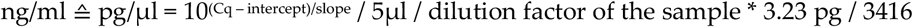

## References

1. Malkin, E.Z.; Bratman, S. V. Bioactive DNA from extracellular vesicles and particles. Cell Death Dis. 2020, 11, 584, doi:10.1038/s41419-020-02803-4.

2. Contrepois, K.; Wu, S.; Moneghetti, K.J.; Hornburg, D.; Ahadi, S.; Tsai, M.S.; Metwally, A.A.; Wei, E.; Lee-McMullen, B.; Quijada, J. V.; et al. Molecular Choreography of Acute Exercise. Cell 2020, doi:10.1016/j.cell.2020.04.043.

3. Uhlén, M.; Karlsson, M.J.; Hober, A.; Svensson, A.-S.; Scheffel, J.; Kotol, D.; Zhong, W.; Tebani, A.; Strandberg, L.; Edfors, F.; et al. The human secretome. Sci. Signal. 2019, 12, eaaz0274, doi:10.1126/scisignal.aaz0274.

4. Stroun, M.; Anker, P. Nucleic acids spontaneously released by living frog auricles. Biochem. J. 1972, doi:10.1042/bj1280100Pb.

5. Anker, P.; Stroun, M.; Maurice, P.A. Spontaneous Release of DNA by Human Blood Lymphocytes as Shown in an in Vitro System. Cancer Res. 1975.

6. Bronkhorst, A.J.; Wentzel, J.F.; Aucamp, J.; van Dyk, E.; du Plessis, L.; Pretorius, P.J. Characterization of the cell-free DNA released by cultured cancer cells. Biochim. Biophys. Acta - Mol. Cell Res. 2016, doi:10.1016/j.bbamcr.2015.10.022.

7. Galvanin, A.; Dostert, G.; Ayadi, L.; Marchand, V.; Velot, É.; Motorin, Y. Diversity and heterogeneity of extracellular RNA in human plasma. Biochimie 2019, 164, 22–36, doi:10.1016/j.biochi.2019.05.011.

8. Beiter, T.; Fragasso, A.; Hudemann, J.; Nieß, A.M.; Simon, P. Short-Term Treadmill Running as a Model for Studying Cell-Free DNA Kinetics In Vivo. Clin. Chem. 2011, 57, 633–636, doi:10.1373/clinchem.2010.158030.

9. Breitbach, S.; Sterzing, B.; Magallanes, C.; Tug, S.; Simon, P. Direct measurement of cell-free DNA from serially collected capillary plasma during incremental exercise. J. Appl. Physiol. 2014, doi:10.1152/japplphysiol.00002.2014.

10. Shah, R.; Yeri, A.; Das, A.; Courtright-Lim, A.; Ziegler, O.; Gervino, E.; Ocel, J.; Quintero-Pinzon, P.; Wooster, L.; Bailey, C.S.; et al. Small RNA-seq during acute maximal exercise reveal RNAs involved in vascular inflammation and cardiometabolic health: Brief report. Am. J. Physiol. - Hear. Circ. Physiol. 2017, doi:10.1152/ajpheart.00500.2017.

11. Moss, J.; Magenheim, J.; Neiman, D.; Zemmour, H.; Loyfer, N.; Korach, A.; Samet, Y.; Maoz, M.; Druid, H.; Arner, P.; et al. Comprehensive human cell-type methylation atlas reveals origins of circulating cell-free DNA in health and disease. Nat. Commun. 2018, doi:10.1038/s41467-018-07466-6.

12. Lui, Y.Y.N.; Woo, K.-S.; Wang, A.Y.M.; Yeung, C.-K.; Li, P.K.T.; Chau, E.; Ruygrok, P.; Lo, Y.M.D. Origin of Plasma Cell-free DNA after Solid Organ Transplantation. Clin. Chem. 2003, 49, 495–496, doi:10.1373/49.3.495.

13. Tug, S.; Helmig, S.; Deichmann, E.R.; Schmeier-Jürchott, A.; Wagner, E.; Zimmermann, T.; Radsak, M.; Giacca, M.; Simon, P. Exercise-induced increases in cell free DNA in human plasma originate predominantly from cells of the haematopoietic lineage. Exerc. Immunol. Rev. 2015.

14. Jiang, P.; Lo, Y.M.D. The Long and Short of Circulating Cell-Free DNA and the Ins and Outs of Molecular Diagnostics. Trends Genet. 2016, 32, 360–371, doi:10.1016/j.tig.2016.03.009.

15. Grabuschnig, S.; Bronkhorst, A.J.; Holdenrieder, S.; Rosales Rodriguez, I.; Schliep, K.P.; Schwendenwein, D.; Ungerer, V.; Sensen, C.W. Putative Origins of Cell-Free DNA in Humans: A Review of Active and Passive Nucleic Acid Release Mechanisms. Int. J. Mol. Sci. 2020, 21, 8062, doi:10.3390/ijms21218062.

16. Yáñez-Mó, M.; Siljander, P.R.M.; Andreu, Z.; Bedina Zavec, A.; Borràs, F.E.; Buzas, E.I.; Buzas, K.; Casal, E.; Cappello, F.; Carvalho, J.; et al. Biological properties of extracellular vesicles and their physiological functions. J. Extracell. Vesicles 2015, 4, 27066, doi:10.3402/jev.v4.27066.

17. Brahmer, A.; Neuberger, E.; Esch-Heisser, L.; Haller, N.; Jorgensen, M.M.; Baek, R.; Möbius, W.; Simon, P.; Krämer-Albers, E.-M. Platelets, endothelial cells and leukocytes contribute to the exercise-triggered release of extracellular vesicles into the circulation. J. Extracell. Vesicles 2019, 8, 1615820, doi:10.1080/20013078.2019.1615820.

18. Frühbeis, C.; Helmig, S.; Tug, S.; Simon, P.; Krämer-Albers, E.-M. Physical exercise induces rapid release of small extracellular vesicles into the circulation. J. Extracell. Vesicles 2015, 4, 28239, doi:10.3402/jev.v4.28239.

19. Whitham, M.; Parker, B.L.; Friedrichsen, M.; Hingst, J.R.; Hjorth, M.; Hughes, W.E.; Egan, C.L.; Cron, L.; Watt, K.I.; Kuchel, R.P.; et al. Extracellular Vesicles Provide a Means for Tissue Crosstalk during Exercise. Cell Metab. 2018, 27, 237–251.e4, doi:10.1016/j.cmet.2017.12.001.

20. Arraud, N.; Linares, R.; Tan, S.; Gounou, C.; Pasquet, J.M.; Mornet, S.; Brisson, A.R. Extracellular vesicles from blood plasma: Determination of their morphology, size, phenotype and concentration. J. Thromb. Haemost. 2014, doi:10.1111/jth.12554.

21. Berckmans, R.J.; Lacroix, R.; Hau, C.M.; Sturk, A.; Nieuwland, R. Extracellular vesicles and coagulation in blood from healthy humans revisited. J. Extracell. Vesicles 2019, doi:10.1080/20013078.2019.1688936.

22. Palviainen, M.; Saraswat, M.; Varga, Z.; Kitka, D.; Neuvonen, M.; Puhka, M.; Joenväärä, S.; Renkonen, R.; Nieuwland, R.; Takatalo, M.; et al. Extracellular vesicles from human plasma and serum are carriers of extravesicular cargo—Implications for biomarker discovery. PLoS One 2020, 15, e0236439, doi:10.1371/journal.pone.0236439.

23. Buzás, E.I.; Tóth, E.Á.; Sódar, B.W.; Szabó-Taylor, K.É. Molecular interactions at the surface of extracellular vesicles. Semin. Immunopathol. 2018, 40, 453–464, doi:10.1007/s00281-018-0682-0.

24. Kalluri, R.; LeBleu, V.S. Discovery of Double-Stranded Genomic DNA in Circulating Exosomes. Cold Spring Harb. Symp. Quant. Biol. 2016, 81, 275–280, doi:10.1101/sqb.2016.81.030932.

25. Thakur, B.K.; Zhang, H.; Becker, A.; Matei, I.; Huang, Y.; Costa-Silva, B.; Zheng, Y.; Hoshino, A.; Brazier, H.; Xiang, J.; et al. Double-stranded DNA in exosomes: a novel biomarker in cancer detection. Cell Res. 2014, 24, 766–769, doi:10.1038/cr.2014.44.

26. Jeppesen, D.K.; Fenix, A.M.; Franklin, J.L.; Higginbotham, J.N.; Zhang, Q.; Zimmerman, L.J.; Liebler, D.C.; Ping, J.; Liu, Q.; Evans, R.; et al. Reassessment of Exosome Composition. Cell 2019, doi:10.1016/j.cell.2019.02.029.

27. Lee, T.H.; Chennakrishnaiah, S.; Audemard, E.; Montermini, L.; Meehan, B.; Rak, J. Oncogenic ras-driven cancer cell vesiculation leads to emission of double-stranded DNA capable of interacting with target cells. Biochem. Biophys. Res. Commun. 2014, doi:10.1016/j.bbrc.2014.07.109.

28. Fischer, S.; Cornils, K.; Speiseder, T.; Badbaran, A.; Reimer, R.; Indenbirken, D.; Grundhoff, A.; Brunswig-Spickenheier, B.; Alawi, M.; Lange, C. Indication of Horizontal DNA Gene Transfer by Extracellular Vesicles. PLoS One 2016, 11, e0163665, doi:10.1371/journal.pone.0163665.

29. Balaj, L.; Lessard, R.; Dai, L.; Cho, Y.J.; Pomeroy, S.L.; Breakefield, X.O.; Skog, J. Tumour microvesicles contain retrotransposon elements and amplified oncogene sequences. Nat. Commun. 2011, doi:10.1038/ncomms1180.

30. Choi, D.; Montermini, L.; Jeong, H.; Sharma, S.; Meehan, B.; Rak, J. Mapping Subpopulations of Cancer Cell-Derived Extracellular Vesicles and Particles by Nano-Flow Cytometry. ACS Nano 2019, doi:10.1021/acsnano.9b04480.

31. Németh, A.; Orgovan, N.; Sódar, B.W.; Osteikoetxea, X.; Pálóczi, K.; Szabó-Taylor, K.É.; Vukman, K. V.; Kittel, Á.; Turiák, L.; Wiener, Z.; et al. Antibiotic-induced release of small extracellular vesicles (exosomes) with surface-associated DNA. Sci. Rep. 2017, 7, 8202, doi:10.1038/s41598-017-08392-1.

32. Zhang, H.; Freitas, D.; Kim, H.S.; Fabijanic, K.; Li, Z.; Chen, H.; Mark, M.T.; Molina, H.; Martin, A.B.; Bojmar, L.; et al. Identification of distinct nanoparticles and subsets of extracellular vesicles by asymmetric flow field-flow fractionation. Nat. Cell Biol. 2018, 20, 332–343, doi:10.1038/s41556-018-0040-4.

33. Zhang, Q.; Higginbotham, J.N.; Jeppesen, D.K.; Yang, Y.-P.; Li, W.; McKinley, E.T.; Graves-Deal, R.; Ping, J.; Britain, C.M.; Dorsett, K.A.; et al. Transfer of Functional Cargo in Exomeres. Cell Rep. 2019, 27, 940–954.e6, doi:10.1016/j.celrep.2019.01.009.

34. Takahashi, A.; Okada, R.; Nagao, K.; Kawamata, Y.; Hanyu, A.; Yoshimoto, S.; Takasugi, M.; Watanabe, S.; Kanemaki, M.T.; Obuse, C.; et al. Erratum to: Exosomes maintain cellular homeostasis by excreting harmful DNA from cells (Nature Communications, (2017), 8, (15287), 10.1038/ncomms15287). Nat. Commun. 2018.

35. Allenson, K.; Castillo, J.; San Lucas, F.A.; Scelo, G.; Kim, D.U.; Bernard, V.; Davis, G.; Kumar, T.; Katz, M.; Overman, M.J.; et al. High prevalence of mutant KRAS in circulating exosome-derived DNA from early-stage pancreatic cancer patients. Ann. Oncol. 2017, doi:10.1093/annonc/mdx004.

36. Kahlert, C.; Melo, S.A.; Protopopov, A.; Tang, J.; Seth, S.; Koch, M.; Zhang, J.; Weitz, J.; Chin, L.; Futreal, A.; et al. Identification of Double-stranded Genomic DNA Spanning All Chromosomes with Mutated KRAS and p53 DNA in the Serum Exosomes of Patients with Pancreatic Cancer. J. Biol. Chem. 2014, 289, 3869–3875, doi:10.1074/jbc.C113.532267.

37. Lázaro-Ibáñez, E.; Lässer, C.; Shelke, G.V.; Crescitelli, R.; Jang, S.C.; Cvjetkovic, A.; García-Rodríguez, A.; Lötvall, J. DNA analysis of low- and high-density fractions defines heterogeneous subpopulations of small extracellular vesicles based on their DNA cargo and topology. J. Extracell. Vesicles 2019, doi:10.1080/20013078.2019.1656993.

38. Yang, S.; Che, S.P.Y.; Kurywchak, P.; Tavormina, J.L.; Gansmo, L.B.; Correa de Sampaio, P.; Tachezy, M.; Bockhorn, M.; Gebauer, F.; Haltom, A.R.; et al. Detection of mutant KRAS and TP53 DNA in circulating exosomes from healthy individuals and patients with pancreatic cancer. Cancer Biol. Ther. 2017, 18, 158–165, doi:10.1080/15384047.2017.1281499.

39. Vagner, T.; Spinelli, C.; Minciacchi, V.R.; Balaj, L.; Zandian, M.; Conley, A.; Zijlstra, A.; Freeman, M.R.; Demichelis, F.; De, S.; et al. Large extracellular vesicles carry most of the tumour DNA circulating in prostate cancer patient plasma. J. Extracell. Vesicles 2018, doi:10.1080/20013078.2018.1505403.

40. Klump, J.; Phillipp, U.; Follo, M.; Eremin, A.; Lehmann, H.; Nestel, S.; von Bubnoff, N.; Nazarenko, I. Extracellular vesicles or free circulating DNA: where to search for BRAF and cKIT mutations? Nanomedicine Nanotechnology, Biol. Med. 2018, doi:10.1016/j.nano.2017.12.009.

41. Helmig, S.; Frühbeis, C.; Krämer-Albers, E.-M.; Simon, P.; Tug, S. Release of bulk cell free DNA during physical exercise occurs independent of extracellular vesicles. Eur. J. Appl. Physiol. 2015, 115, 2271–2280, doi:10.1007/s00421-015-3207-8.

42. Cai, J.; Guan, W.; Tan, X.; Chen, C.; Li, L.; Wang, N.; Zou, X.; Zhou, F.; Wang, J.; Pei, F.; et al. SRY gene transferred by extracellular vesicles accelerates atherosclerosis by promotion of leucocyte adherence to endothelial cells. Clin. Sci. 2015, doi:10.1042/CS20140826.

43. Fernando, M.R.; Jiang, C.; Krzyzanowski, G.D.; Ryan, W.L. New evidence that a large proportion of human blood plasma cell-free DNA is localized in exosomes. PLoS One 2017, 12, e0183915, doi:10.1371/journal.pone.0183915.

44. Monguió-Tortajada, M.; Gálvez-Montón, C.; Bayes-Genis, A.; Roura, S.; Borràs, F.E. Extracellular vesicle isolation methods: rising impact of size-exclusion chromatography. Cell. Mol. Life Sci. 2019, 76, 2369–2382, doi:10.1007/s00018-019-03071-y.

45. Simonsen, J.B. What Are We Looking At? Extracellular Vesicles, Lipoproteins, or Both? Circ. Res. 2017, 121, 920–922, doi:10.1161/CIRCRESAHA.117.311767.

46. Jeppesen, D.K.; Fenix, A.M.; Franklin, J.L.; Higginbotham, J.N.; Zhang, Q.; Zimmerman, L.J.; Liebler, D.C.; Ping, J.; Liu, Q.; Evans, R.; et al. Reassessment of Exosome Composition. Cell 2019, 177, 428–445.e18, doi:10.1016/j.cell.2019.02.029.

47. Greening, D.W.; Xu, R.; Ji, H.; Tauro, B.J.; Simpson, R.J. A Protocol for Exosome Isolation and Characterization: Evaluation of Ultracentrifugation, Density-Gradient Separation, and Immunoaffinity Capture Methods. In Methods in Molecular Biology; 2015; pp. 179–209.

48. Brahmer, A.; Neuberger, E.W.I.; Simon, P.; Krämer-Albers, E.-M. Considerations for the Analysis of Small Extracellular Vesicles in Physical Exercise. Front. Physiol. 2020, 11, 1611, doi:10.3389/fphys.2020.576150.

49. Ritchie, C. Rating of Perceived Exertion (RPE). J. Physiother. 2012, 58, 62, doi:10.1016/S1836-9553(12)70078-4.

50. Lacroix, R.; Judicone, C.; Poncelet, P.; Robert, S.; Arnaud, L.; Sampol, J.; Dignat-George, F. Impact of pre-analytical parameters on the measurement of circulating microparticles: towards standardization of protocol. J. Thromb. Haemost. 2012, 10, 437–46, doi:10.1111/j.1538-7836.2011.04610.x.

51. Van Deun, J.; Mestdagh, P.; Agostinis, P.; Akay, Ö.; Anand, S.; Anckaert, J.; Martinez, Z.A.; Baetens, T.; Beghein, E.; Bertier, L.; et al. EV-TRACK: Transparent reporting and centralizing knowledge in extracellular vesicle research. Nat. Methods 2017.

52. Neuberger, E.W.I.; Brahmer, A.; Ehlert, T.; Kluge, K.; Philippi, K.F.A.; Boedecker, S.C., Weinmann-Menke, J.; Simon, P. Validating quantitative PCR assays for cell-free DNA detection without DNA extraction: Exercise induced kinetics in systemic lupus erythematosus patients. doi: https://doi.org/10.1101/2021.01.17.21249972.

53. Sumenkova, D. V; Polyakov, L.M.; Panin, L.E. Plasma Lipoproteins as a Transport Form of Extracellular DNA. Bull. Exp. Biol. Med. 2013, 154, 622–623, doi:10.1007/s10517-013-2014-7.

54. Breitbach, S.; Sterzing, B.; Magallanes, C.; Tug, S.; Simon, P. Direct measurement of cell-free DNA from serially collected capillary plasma during incremental exercise. J. Appl. Physiol. 2014, 117, 119–130, doi:10.1152/japplphysiol.00002.2014.

55. Tug, S.; Mehdorn, M.; Helmig, S.; Breitbach, S.; Ehlert, T.; Simon, P. Exploring the Potential of Cell-Free-DNA Measurements After an Exhaustive Cycle-Ergometer Test as a Marker for Performance-Related Parameters. Int. J. Sports Physiol. Perform. 2017, 12, 597–604, doi:10.1123/ijspp.2016-0157.

56. Napirei, M.; Wulf, S.; Eulitz, D.; Mannherz, H.G.; Kloeckl, T. Comparative characterization of rat deoxyribonuclease 1 (Dnase1) and murine deoxyribonuclease 1-like 3 (Dnase1l3). Biochem. J. 2005, 389, 355–64, doi:10.1042/BJ20042124.

57. Bronkhorst, A.J.; Ungerer, V.; Holdenrieder, S. The emerging role of cell-free DNA as a molecular marker for cancer management. Biomol. Detect. Quantif. 2019, 17, 100087, doi:10.1016/j.bdq.2019.100087.

58. Möller, A.; Lobb, R.J. The evolving translational potential of small extracellular vesicles in cancer. Nat. Rev. Cancer 2020.

59. Kowal, J.; Arras, G.; Colombo, M.; Jouve, M.; Morath, J.P.; Primdal-Bengtson, B.; Dingli, F.; Loew, D.; Tkach, M.; Théry, C. Proteomic comparison defines novel markers to characterize heterogeneous populations of extracellular vesicle subtypes. Proc. Natl. Acad. Sci. U. S. A. 2016, doi:10.1073/pnas.1521230113.

60. Van Deun, J.; Mestdagh, P.; Sormunen, R.; Cocquyt, V.; Vermaelen, K.; Vandesompele, J.; Bracke, M.; De Wever, O.; Hendrix, A. The impact of disparate isolation methods for extracellular vesicles on downstream RNA profiling. J. Extracell. Vesicles 2014, doi:10.3402/jev.v3.24858.

61. Lobb, R.J.; Becker, M.; Wen, S.W.; Wong, C.S.F.; Wiegmans, A.P.; Leimgruber, A.; Möller, A. Optimized exosome isolation protocol for cell culture supernatant and human plasma. J. Extracell. Vesicles 2015, doi:10.3402/jev.v4.27031.

62. Tandia, B.-M.; Vandenbranden, M.; Wattiez, R.; Lakhdar, Z.; Ruysschaert, J.-M.; Elouahabi, A. Identification of human plasma proteins that bind to cationic lipid/DNA complex and analysis of their effects on transfection efficiency: implications for intravenous gene transfer. Mol. Ther. 2003, 8, 264–273, doi:10.1016/S1525-0016(03)00150-3.

63. Qubit^®^ dsDNA HS Assay Kits. Available online: https://www.thermofisher.com/document-connect/document-connect.html?url= https%3A%2F%2Fassets.thermofisher.com%2FTFS-Assets%2FLSG%2Fmanuals%2FQubit_dsDNA_HS_Assay_UG.pdf&title=VXNlciBHdWlkZTogUXViaXQgZHNETkEgSFMgQXNzYXkgS2l0cw== (accessed on Jan 27, 2021).

64. Vechetti, I.J.; Valentino, T.; Mobley, C.B.; McCarthy, J.J. The role of extracellular vesicles in skeletal muscle and systematic adaptation to exercise. J. Physiol. 2020.

65. Fuller, O.K.; Whitham, M.; Mathivanan, S.; Febbraio, M.A. The Protective Effect of Exercise in Neurodegenerative Diseases: The Potential Role of Extracellular Vesicles. Cells 2020, 9, 2182, doi:10.3390/cells9102182.

66. Denham, J.; Spencer, S.J. Emerging roles of extracellular vesicles in the intercellular communication for exercise-induced adaptations. Am. J. Physiol. Metab. 2020, 319, E320–E329, doi:10.1152/ajpendo.00215.2020.

